# Lactate production can function to increase human epithelial cell iron concentration

**DOI:** 10.1101/2021.11.11.468295

**Authors:** Caroline Ghio, Joleen M. Soukup, Lisa A. Dailey, Andrew J. Ghio, Dina M. Schreinemachers, Ryan A. Koppes, Abigail N. Koppes

## Abstract

Iron is an essential micronutrient required by every cell, inclusive of both prokaryotes and eukaryotes. Under conditions of limited iron availability, plants and microbes evolved mechanisms to acquire iron which include carbon metabolism reprogramming, with the activity of several enzymes involved in the Krebs cycle and the glycolytic pathway being stimulated by metal deficiency. Following release, resultant carboxylates/hydroxycarboxylates can function as ligands to complex iron and facilitate its solubilization and uptake, reversing the deficiency. Human epithelial tissue may produce lactate, a hydroxycarboxylate, during absolute and functional iron deficiency in an attempt to import metal to reverse limited availability. Here we investigate 1) if lactate can increase cell metal import, 2) if lactic dehydrogenase (LDH) activity in and lactate production by cells correspond to metal availability, and 3) if blood concentrations of LDH in a human cohort correlate with indices of iron homeostasis. Exposures of Caco-2 cells to both Na lactate and ferric ammonium citrate (FAC) increased metal import relative to FAC alone. Fumaric, isocitric, malic, and succinic acid exposure revealed that FAC co-incubation similarly increased iron import relative to FAC alone. Increased iron import following exposures to Na lactate and FAC elevated both ferritin and metal associated with mitochondria. LDH in Caco-2 cell scrapings did not change after exposure to deferoxamine but decreased with 24 hr exposure to FAC. Lactate levels in both the supernatants and cell scrapings revealed decreased levels at 4, 8, and 24 hr with FAC. In the National Health and Nutrition Examination Survey (NHANES 2005-2010), Spearman correlations demonstrated significant negative relationships between LDH concentrations and serum iron. We conclude that iron import in human cells can involve lactate, LDH activity can reflect the availability of this metal, and blood LDH concentrations can correlate with indices of iron homeostasis.

## Introduction

A favorable oxidation-reduction potential and an environmental abundance has led to the evolutionary selection of iron for a wide range of fundamental functions in cells ^1–3^. As a result, life is ferrocentric with iron being an essential micronutrient required by almost every living system. However, the limited amount of ferric ion (Fe^3+^) available on Earth meant that human survival required adaptations to produce greater quantities of metal to support life ^1–3^. This challenge was realized by cells and systems utilizing combinations of two pathways to acquire essential iron: 1) the chemical reduction of Fe^3+^ to Fe^2+^ (i.e. ferri-reduction) with subsequent import and 2) the complexation of Fe^3+^ with chelators coupled with receptors for uptake. In addition to solubility limiting its availability, iron-catalyzed generation of radicals presented the potential for oxidative stress. Such reactivity mandated that iron homeostasis be tightly controlled. Therefore, the import, storage, and efflux of this metal are carefully regulated, and life is positioned at the interface between iron deficiency and -sufficiency ^1–3^.

To better understand the mechanisms of iron homeostasis, significant work has centered on the intestinal lining, where most dietary iron is absorbed. Due to the complexities of whole organism investigation, *in vitro* models are paramount to understanding cellular regulation. A human epithelial cell line, Caco-2 have been demonstrated to be an effective cell line in modeling iron homeostasis in the intestinal lining. Caco-2 cells respond to treatment with iron through incubation with Fe-NTA and measuring ferritin, an iron storage protein, transferrin receptor, a glycoprotein that mediates cellular iron uptake, DMT1, an iron uptake transporter, and ferroportin 1, an iron export transporter ^4^. When cells are exposed to iron, DMT1, transferrin receptor, and ferroportin 1 levels decrease, while ferritin levels increase ^4^. In Caco-2 cells researchers have also found five distinct gene clusters with unique expression profiles according to iron status which suggests that iron metabolism may affect immune response and lipid metabolism ^5^. Additionally, an increase in intracellular iron has also been shown to increase calreticulin, a chaperone protein that aids in the folding of proteins synthesized in the endoplasmic reticulum ^6^. The balance in iron homeostasis and the importance of iron sufficiency are clearly seen in these human epithelial cultures, however the underlying mechanisms of iron import remains unknown.

Mechanisms to acquire iron under conditions of limited availability, which evolved in plants and microbes, can include a reprogramming of carbon metabolism with the activity of several enzymes involved in the Krebs cycle and the glycolytic pathway being stimulated following metal deficiency ^7^. Production and release of carboxylates/hydroxycarboxylates can result in these compounds demonstrating an affinity for metal complexation ^8,9^. After their secretion, carboxylates/hydroxycarboxylates function as ligands to complex iron facilitating solubilization, reduction, and uptake of the metal ^10^. In plants, the total concentration of carboxylates/hydroxycarboxylates (e.g. malate and citrate) increases in xylem and both root and leaf extracts during iron deficiency ^11^. Yeasts similarly increase the production of carboxylates/hydroxycarboxylates (i.e. malate, succinate, and fumarate) in response to iron-deficiency ^12^.

Through glycolysis, glucose is metabolized to pyruvate which is oxidized to water and carbon dioxide through the Krebs cycle and oxidative phosphorylation in the mitochondria. If oxidation is not possible as a result of a deficiency in required substrates (e.g. oxygen), pyruvate is converted into lactate through lactic dehydrogenase (LDH) with concomitant interconversion of nicotinamide adenine dinucleotide in reduced and oxidized form (NADH and NAD^+^ respectively). The expression of LDH can be impacted by iron deficiency in both plants and microbes supporting a role in metal homeostasis ^13–15^. Its product, lactate, is an α-hydroxycarboxylate, which complexes iron comparable to plant and microbial siderophores ^16,17^. Such binding with ensuing transport expedites cell iron import ^18^. It is proposed that comparable to plant cells and microbes, human cells produce lactate during iron deficiency in an attempt to import greater quantities of the metal and reverse its limited availability. In this study, we investigate 1) if lactate can increase the *in vitro* import of iron into human cells, 2) if the *in vitro* activity of LDH within and the concentrations of lactate released by these cells correspond to the availability of the metal, and 3) whether blood concentrations of LDH in a human cohort correlate with indices of iron homeostasis.

## Materials and Methods

### Materials

All reagents were from Sigma (St. Louis, MO) unless specified otherwise. Ferric ammonium citrate (FAC) was used as the *in vitro* source of iron because it is buffered and considered physiologically relevant. The Hanks Buffered Salt Solution (HBSS) used was supplemented with calcium and magnesium.

### Cell cultures

As a result of their frequent employment as a model for *in vitro* studies of iron uptake, human epithelial colorectal adenocarcinoma cells, Caco-2 (HTB-37; ATCC, Manassas, VA), were used in this investigation^19,20^. These were grown in Dulbecco’s modified Eagle’s medium (DMEM, Fisher Scientific) containing 10% fetal bovine serum (FBS; ATCC) and 1% antibiotic-antimitotic solution of 0.25% pen-strep (Fisher Scientific). Unless specified otherwise, Caco-2 cells were seeded in a tissue culture treated 12-well plate and grown to 90–100% confluence by observation (Fisher Scientific). The medium was replaced every other day.

An immortalized line of primary human bronchial epithelium derived by transfection of primary cells with SV40 early-region genes (BEAS-2B) was utilized to identify any differences in human epithelial tissues. BEAS-2B cells were grown to 90-100% confluence on uncoated tissue culture treated plastic 12-well plates in keratinocyte growth medium (Clonetics) which is essentially MCDB 153 medium supplemented with 5 ng/mL human epidermal growth factor, 5 mg/mL insulin, 0.5 mg/mL hydrocortisone, 0.15 mM calcium, bovine pituitary extract, 0.1 mM ethanolamine and 0.1 mM phosphoethanolamine. Unless specified otherwise, BEAS-2B cells were also seeded in a 12-well plate and grown to 90–100% confluence (Fisher Scientific). Fresh medium was provided every other day.

### Cell iron import measurement with ICPOES

Import of iron was examined in response to 4-hr exposures in culture. To examine the uptake of metal lactates, Caco-2 cells were exposed (3 wells per plate each) to a) HBSS, b) 500 μM iron lactate, c) 500 μM aluminum lactate, and d) 500 μM zinc lactate. To delineate differences in the uptake of iron from different compounds, Caco-2 cells were exposed (3 wells per plate each) to a) HBSS, b) 500 μM FAC, c) 500 μM ferric citrate, and d) 500 μM ferrous lactate. To examine for differences in the uptake of iron between ferrous and ferric compounds, Caco-2 cells were exposed (4 wells per plate each) to a) HBSS, b) 500 μM ferrous sulfate, and c) 500 μM ferric sulfate for 4 hr. To examine the effect of lactate on iron import, Caco-2 cells were grown in 12 well plates and exposed (3 wells per plate each) to a) HBSS, b) 200 μM FAC, c) 0.5 and 5.0 mM sodium (Na) lactate, and d) both 200 μM FAC and 0.5 and 5.0 mM Na lactate. To evaluate the influence of FAC and sodium lactate, Caco-2 cells were grown in 12 well plates and exposed (3 wells per plate each) to a) HBSS, b) 200 μM FAC, c) 0.5 and 5.0 mM lactic acid, and d) both 200 μM FAC and 0.5 and 5.0 mM lactic acid. Caco-2 cells were exposed to 200 μM FAC both with and without 500 μM fumaric acid, isocitric acid, malic acid, and succinic acid.

To evaluate if the tissue of origin influenced cell iron import, BEAS-2B cells were grown in 12 well plates and exposed (3 wells per plate each) to a) HBSS, b) 200 μM FAC, c) 0.5 and 5.0 mM Na lactate, and d) both 200 μM FAC and 0.5 and 5.0 mM Na lactate. Additionally, BEAS-2B cells were grown in 12 well plates and exposed (3 wells per plate each) to a) HBSS, b) 200 μM FAC, c) 0.5 and 5.0 mM lactic acid, and d) both 200 μM FAC and 0.5 and 5.0 mM lactic acid.

For all experimental conditions, after incubation, the cells were gently washed with HBSS, scraped into 10% trichloroacetic acid (TCA) dissolved in 1.0 mL of 3 N HCl, and digested at 70° C for 24 hr. Concentrations of metals in the cell lysates were determined using inductively coupled plasma optical emission spectroscopy (ICPOES; Model Optima 4300D; Perkin Elmer, Norwalk, CT) operated at wavelengths of 238.204 and 259.939 nm for iron, 396.215 and 308.215 nm for aluminum, and 206.200 and 213.857 nm for zinc.

### Cell ferritin concentrations

Caco-2 and BEAS-2B cells were exposed to a) HBSS, b) 200 μM FAC, c) 500 μM Na lactate, and d) both 200 μM FAC and 500 μM Na lactate for 24 hr. After removal of the media, cells were scraped into 1.0 mL PBS and disrupted using five passes through a small gauge needle. The ferritin concentrations in the lysates were quantified using an immunoturbidimetric assay (Kamiya Biomedical Company).

### Mitochondrial iron concentrations

Caco-2 and BEAS-2B cells were grown in 75-cm^2^ flasks, incubated in a) HBSS, b) 200 μM FAC, c) 500 μM Na lactate, and d) both 200 μM FAC and 500 μM Na lactate for 4 hr. The cells were washed with HBSS and nuclear and mitochondrial fractions were collected using established protocols ^21^. Fractions were digested in 1.0 mL 3 N HCl/10% TCA at 70° C for 24 hr. Non-heme iron concentrations were determined using ICPOES.

### Assay for oxidant production

The potential generation of oxidants by iron with and without lactate was quantified as thiobarbituric acid (TBA) reactive products of deoxyribose. The pentose sugar 2-deoxy-D-ribose reacts with oxidants to yield a mixture of products. On heating with TBA at low pH, these products form a pink chromophore that can be measured by its absorbance at 532 nm. This chromophore is indistinguishable from a TBA-malonaldehyde adduct. The reaction mixture, in a total volume of 2.0 ml of HBSS, contained the following reagents: 1.0 mM deoxyribose, 1.0 mM H_2_O_2_, 1.0 mM ascorbate, and either HBSS or 0.5 mM Fe_2_(SO_4_)_3_. Na lactate and deferoxamine were included at 10 mM. The mixtures were incubated at 37°C for 2 hr with agitation and then centrifuged at 1,200 g for 10 min. One milliliter of both 1.0% (wt/vol) TBA and 2.8% (wt/vol) trichloroacetic acid were added to 1.0 ml of supernatant, heated at 100°C for 10 min, cooled in ice, and the chromophore determined by its absorbance at 532 nm. Measurements were done in replicates of six.

### RT-PCR

Caco-2 cells were exposed to HBSS, deferoxamine in HBSS (50 μM), and FAC in HBSS (500 μM) for 24 hr. Total RNA was isolated using a RNeasy Mini kit (Qiagen, Valencia, CA) and reverse transcribed to generate cDNA using the iScript cDNA Synthesis kit (BioRad, Hercules, CA). Oligonucleotide primer pairs and fluorescent probes for hypoxia-inducible factor −1 (HIF-1α), LDH, and β-Actin were designed using a primer design program (Primer Express; Applied Biosystems) and obtained from Integrated DNA Technologies (Coralville, IA). Quantitative fluorogenic amplification of cDNA was performed using the StepOnePlus Real-Time PCR System (Applied Biosystems), primer/probe sets of interest, and iTaq Universal Probes Supermix (BioRad). The relative abundance of β-Actin mRNA was used to normalize mRNA levels.

### Cell lactate dehydrogenase (LDH) concentrations

Caco-2 cells were exposed to HBSS, deferoxamine in HBSS (25 and 50 μM), and FAC in HBSS (100 and 500 μM) for 4, 8, and 24 hr. Cells were scraped into 1.0 mL HBSS. LDH concentrations were measured using a commercially prepared kit (ThermoTrace LD-L Kit). The assay was modified for automated measurement (KoneLab 30, Espoo, Finland).

### Cell and supernatant lactate concentrations

Caco-2 cells were exposed to HBSS, deferoxamine in HBSS (25 and 50 μM), and FAC in HBSS (100 and 500 μM) for 4, 8, and 24 hr. The supernatant was taken. Cells were scraped into 1.0 mL HBSS. Lactate concentrations in the supernatants and scraped cells were measured using a commercially prepared kit (Roche Diagnostics Corp.). The assay was modified for automated measurement (KoneLab 30).

### Associations between LDH and indices of iron homeostasis in a cohort of healthy subjects

While a publicly accessible database in which the associations of lactate with indices of iron homeostasis was not available, the National Health and Nutrition Examination Survey (NHANES) provided LDH concentrations. Associations of LDH with indices of iron homeostasis were investigated in healthy subjects in the combined, publicly available NHANES 2005-2006, 2007-2008, and 2009-2010 databases (NHANES 2005-2010) ^22–24^. Data for ferritin, a blood protein involved in iron homeostasis, was available only for females. Accordingly, the study was limited to non-pregnant females, age 20-49 years. To obtain a healthy study population that would allow determination of the relationships between LDH and endpoints of iron homeostasis without confounding by disease, medications, or smoking, females previously diagnosed with thyroid disease, prediabetes, diabetes, heart failure, coronary heart disease, stroke, and cancer, as well as females prescribed antineoplastic, cardiovascular, hormonal, or metabolic prescriptions, and females defined as active smokers based on high serum cotinine levels (≥10 ng/mL) were excluded from the study. Data on the following endpoints were collected: LDH, iron, ferritin, hemoglobin (HGB), mean cell hemoglobin (MCH), mean corpuscular volume (MCV), and ferritin. SAS statistical software (v9.4) was used for the analyses, including Spearman correlations.

### Statistical Analysis

Data are expressed as mean values ± standard deviation unless specified otherwise. The minimum number of replicates for RTPCR was four while the minimum number of replicates for all other measurements was six. Differences between multiple groups were compared using one-way analysis of variance (ANOVA); the posthoc test employed was Duncan’s Multiple Range test the posthoc tests employed was Duncan’s Multiple Range test and Tukey’s Multiple Comparison test. Two-tailed tests of significance were employed. Significance was assumed at p<0.05.

## Results

### Cell import of metals is elevated by iron lactate availability and ferric ammonium citrate co-incubation

Human gut epithelial cells possess endogenous levels of iron and zinc when cultured in HBSS (**Figure 1A**). Among exposures to metal lactates, those incubated with iron lactate elevated the cell metal concentration to its greatest level (**Figure 1A**). Both aluminum and zinc lactates significantly increased the cell aluminum and zinc concentrations respectively but to a lesser value relative to iron levels after iron lactate. Exposures to iron compounds demonstrated that both ferric ammonium citrate (FAC) and ferric citrate increased Caco-2 cell iron levels but these were diminished relative to those after iron lactate (**Figure 1B**). Relative to HBSS, exposures of Caco-2 cells exposed to ferric and ferrous sulfates revealed significant elevations in cell iron (0.17 ppm ± 0.01, 0.78 ppm ± 0.06, and 0.71 ppm ± 0.09 respectively at 4 hour and 0.18 ppm ± 0.02, 3.31 ppm ± 0.38, and 3.40 ppm ± 0.28 respectively at 24 hour) but there were no significant differences in cell iron between the two iron sulfates. However, while no change is observed in control conditions between 4 and 24 hr incubations, iron intake in the presence of both sulfates increases at a relatively stable ~0.13 ppm/hr. These data support that iron lactate may be preferred by Caco-2 cells for import relative to other sources of metal.

**Figure 1.**
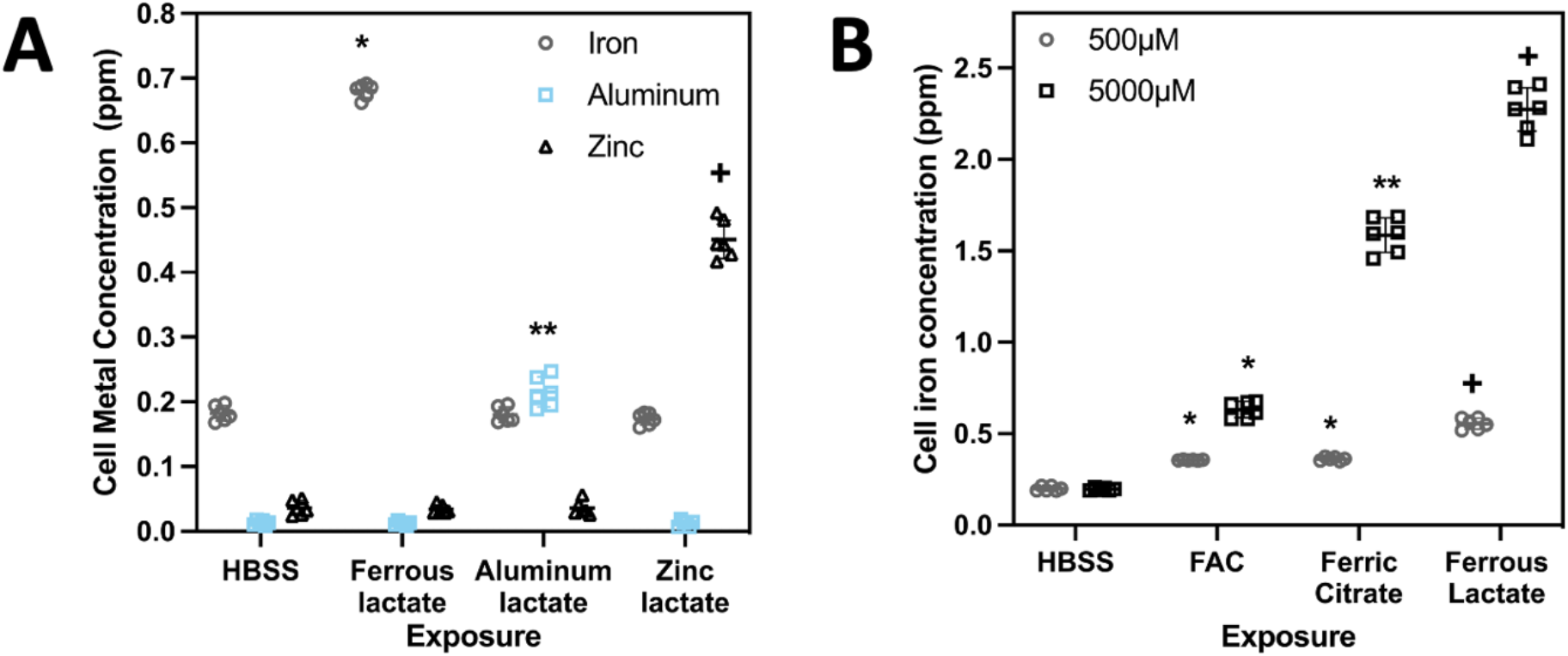
Human epithelial cultures exhibit large amounts of iron import in with the addition of ferrous lactate. Concentrations of Iron, Aluminum, and Zinc differ after Caco-2 cells were exposed to HBSS and 500 μM iron lactate, aluminum lactate, and zinc lactate for 4 hr (A). Increases in imported iron concentration into epithelial cells exposed to iron compounds increased with the addition of ferric ammonium citrate (FAC) and ferric citrate, however the largest import was exhibited with the addition of ferrous lactate for 4 hr (B). However, a 10 fold increase in free iron resulted in 2-fold, 5-fold, and 4-fold increase in imported iron, respectively. * = significantly increased [Fe] relative to HBSS, **= significantly increased [Fe] relative to HBSS and FAC, and + = significantly increased [Fe] relative to all other exposures by ANOVA.

While exposure to Na lactate did not impact Caco-2 cell iron concentrations, incubations with both FAC and Na lactate significantly elevated metal levels relative to FAC alone (0.328 ppm ± 0.022 with FAC alone, 0.455 ppm ± 0.020 with FAC and 0.5 μM Na lactate, 0.334 ppm ± 0.025 with FAC alone, 0.500 ppm ± 0.030 with FAC and 5.0 μM Na lactate) (**Figure 2A**). While the presence of Na lactate increased metal content, the different concentrations (0.5 μM and 5 μM) did exhibit a significant difference in iron intake. BEAS-2B exposures to FAC similarly elevated cell iron while incubations with both FAC and lactate increased metal import to higher levels (0.373 ppm ± 0.020 with FAC alone, 0.551 ppm ± 0.026 with FAC and 0.5 μM Na lactate and 0.382 ppm ± 0.018 with FAC alone, 0.582 ppm ± 0.026 with FAC and 5 μM Na lactate) (**Figure 2B**). Comparable to Na lactate, exposures to both FAC and lactic acid in both Caco-2 and BEAS-2B resulted in increased iron concentrations relative to FAC alone (Caco-2 cells: 0.337 ppm ± 0.020 with FAC, 0.435 ppm ± 0.015 with FAC and 0.5 μM lactic acid and 0.345 ppm ± 0.017 with FAC, 0.444 ppm ± 0.022 with FAC and 5 μM lactic acid) (BEAS-2B cells: 0.384 ppm ± 0.011 with FAC alone, 0.483 ppm ± 0.020 with FAC and 0.5 μM lactic acid and 0.383 ppm ± 0.013 with FAC, 0.464 ppm ± 0.030 p = 0.0009 with FAC and 5 μM lactic acid) (**Figures 2C and 2D**).

**Figure 2.**
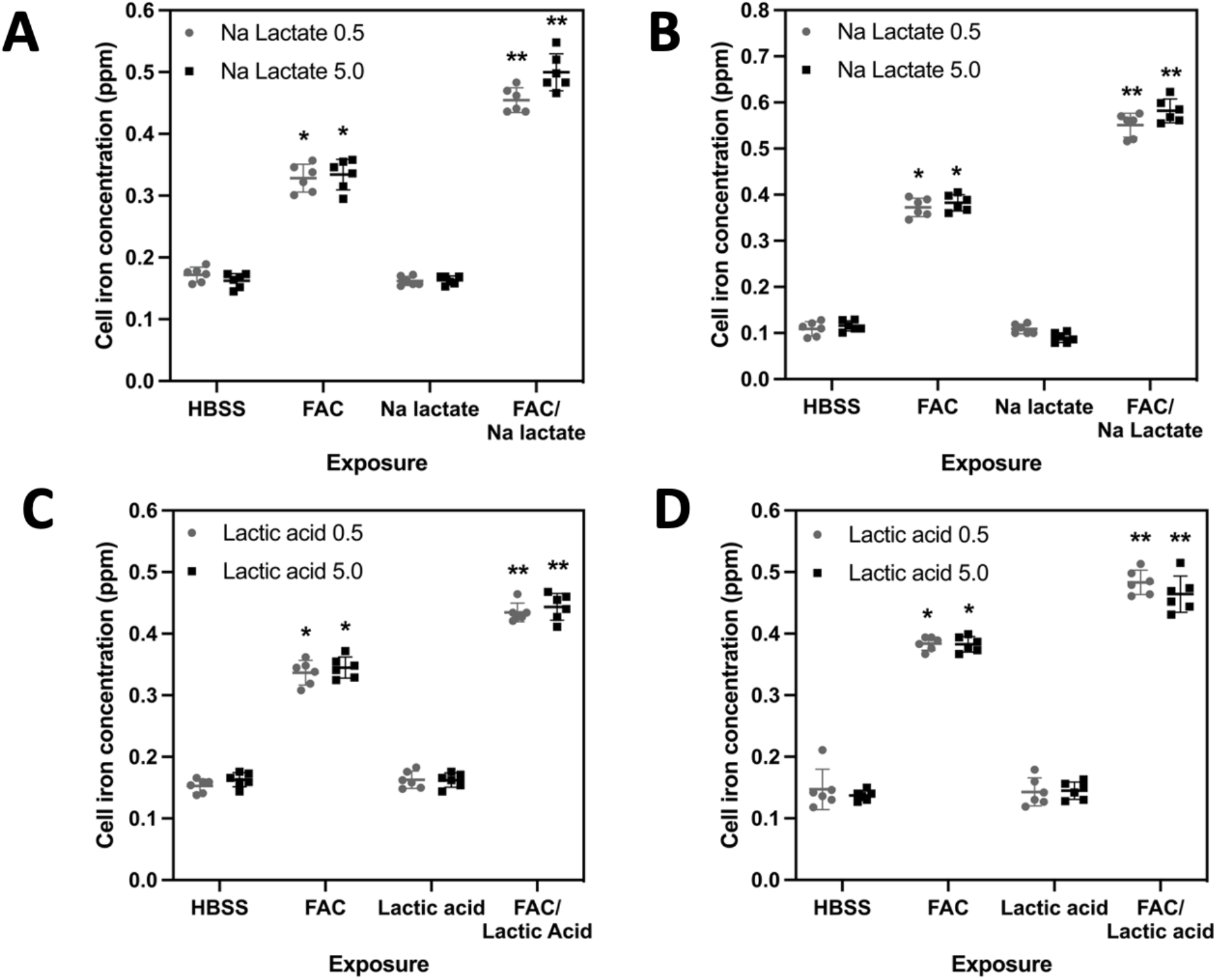
Epithelial cell iron import after exposures to FAC, sodium lactate, and lactic acid indicates no difference across two different tissue sources, and the significant role FAC plays on measured iron. Differences between [Fe] after Caco-2 and BEAS-2B cells were exposed to HBSS, FAC, Na lactate, and co-incubated with FAC and Na lactate for 4 hr (A, B). Differences between [Fe] are apparent after Caco-2 cells and BEAS-2B were exposed to HBSS, FAC, lactate acid, and both FAC and lactate acid for 4 hr (C, D). *= significantly increased [Fe] relative to HBSS, **= significantly increased [Fe] relative to all other exposures; ANOVA.

A dose-response of cell iron with increasing lactic acid was not observed. Exposures of Caco-2 cells with fumaric, isocitric, malic, and succinic acids revealed that their co-incubation with FAC significantly increased import of iron relative to that following FAC alone (0.270 ppm ± 0.023 with FAC alone, 0.401 ppm ± 0.022 with FAC and fumaric acid; 0.307 ppm ± 0.025 with FAC alone, 0.680 ppm ± 0.031 with FAC and isocitric acid; 0.289 ppm ± 0.019 with FAC alone, 0.655 ppm ± 0.023 with FAC and malic acid; 0.281 ppm ± 0.016 with FAC alone, 0.376 ppm ± 0.019 with FAC and succinic acid; **Figures 3A-3D**). The hydroxycarboxylates (i.e. isocitric and malic acids) appeared to double cell iron relative to the dicarboxylates (i.e. fumaric and succinic acids) that only exhibit a one-third increase over FAC exposure alone.

**Figure 3.**
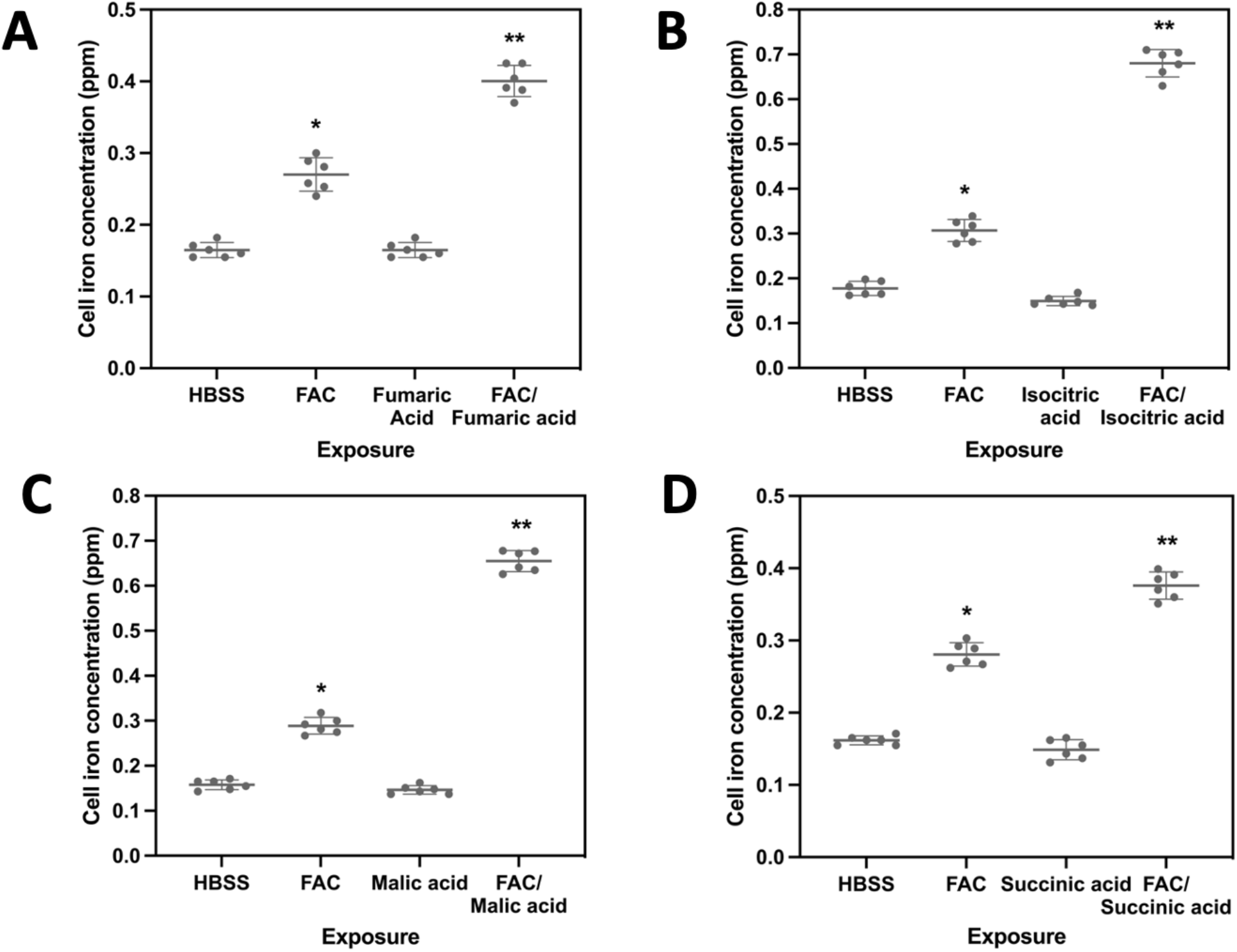
Cell iron import after exposures to FAC and α-hydroxycarboxylic and polycarboxylic acids other than lactate. Ferric ammonia citrate (FAC) increases the iron intake in the presence in both α-hydroxycarboxylic and polycarboxylic acids. Differences in [Fe] after Caco-2 cells were exposed to HBSS, FAC, carboxylate, and both FAC and carboxylate for 4 hr with fumaric acid (A), isocitric acid (B), malic acid (C), and succinic acid (D). *= significantly increased [Fe] relative to HBSS, **= significantly increased [Fe] relative to all other exposures.

Exposures of Caco-2 and BEAS-2B cells to FAC and both FAC and Na lactate impacted cell ferritin levels (37.833 ppm ± 6.676 with FAC alone, 63.833 ppm ± 13.378 with FAC and Na lactate in Caco-2 cells; 47.667 ppm ± 8.238 with FAC alone, 67.833 ppm ± 11.822 with FAC and Na lactate in BEAS-2B cells) (**Figures 4A and 4B**). Relative to FAC alone, cell concentrations of this metal-storage protein were significantly elevated following incubations of both FAC and Na lactate in both cell types. After comparable exposures of Caco-2 and BEAS-2B cells, isolated mitochondria showed an increased iron concentration following incubation with FAC (0.049 ppm ± 0.006 with FAC alone, 0.069 ppm ± 0.008 with FAC and Na lactate in nuclear fractions in Caco-2 cells; 0.044 ppm ± 0.006 with FAC alone, 0.066 ppm ± 0.014 with FAC and Na lactate in mitochondrial fractions in Caco-2 cells; 0.053 ppm ± 0.011 with FAC alone, 0.083 ppm ± 0.009 with FAC and Na lactate in nuclear fractions in BEAS-2B cells; 0.045 ppm ± 0.006 with FAC alone, 0.072 ppm ± 0.009 with FAC and Na lactate in mitochondrial fractions; **Figures 4C and 4D**). With exposure to both FAC and Na lactate, iron levels in the mitochondria were elevated 50% over controls. Both FAC and Na lactate increase iron levels with increased cell expression of ferritin, which is dependent on iron availability, and elevated mitochondrial concentrations.

**Figure 4.**
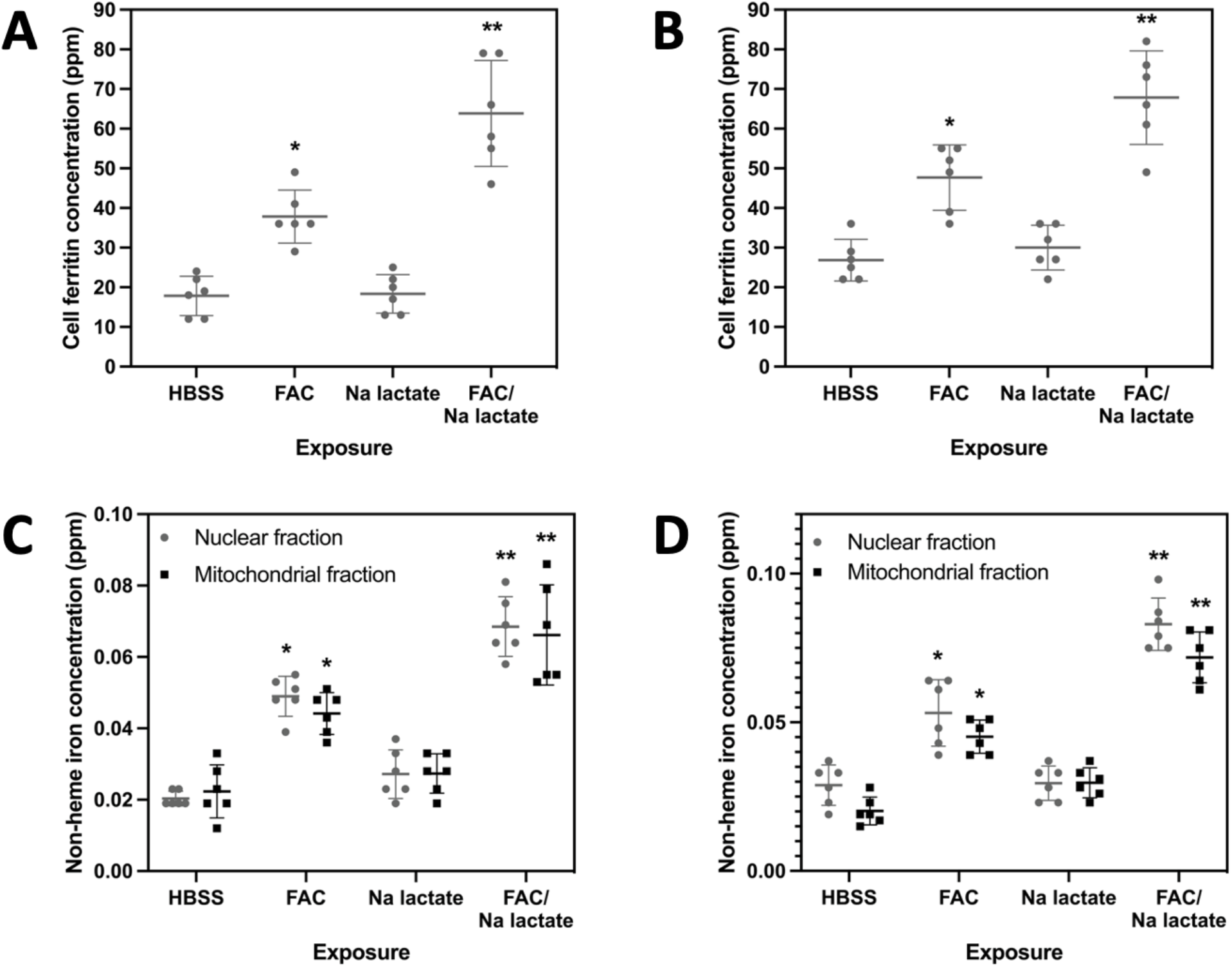
Cell ferritin and mitochondrial iron concentrations. Differences in ferritin concentrations after Caco-2 and BEAS-2B cells were exposed to HBSS, FAC, Na lactate, and both FAC and Na lactate for 24 hr (A and B respectively). Similarly, [Fe] differs in nuclear and mitochondrial fractions after Caco-2 and BEAS-2B cells were exposed to HBSS, FAC, Na lactate, and both FAC and Na lactate for 4 hr (C and D). For A, B, C, and D, *= significantly increased relative to HBSS, **= significantly increased relative to all other exposures, p<0.05.

### Lactate and iron affect absorbance of thiobarbituric acid (TBA) reactive products

Iron may increase oxidizer generation, and this was quantified using TBA reactive products. In an acellular incubation, Fe3+ increased oxidant generation (0.049 ± 0.002 with HBSS, 0.261 ± 0.008 with Fe3+) (**Figure 5**). However, the addition of lactate and deferoxamine significantly decreased the absorbance of TBA reactive products of deoxyribose (0.261 ± 0.008 with HBSS, 0.062 ± 0.003 with lactate, 0.032 ± 0.001 with deferoxamine) (**Figure 5**). This supports the transport of iron by lactate in a coordination complex, which acts to control potential oxidant generation.

**Figure 5.**
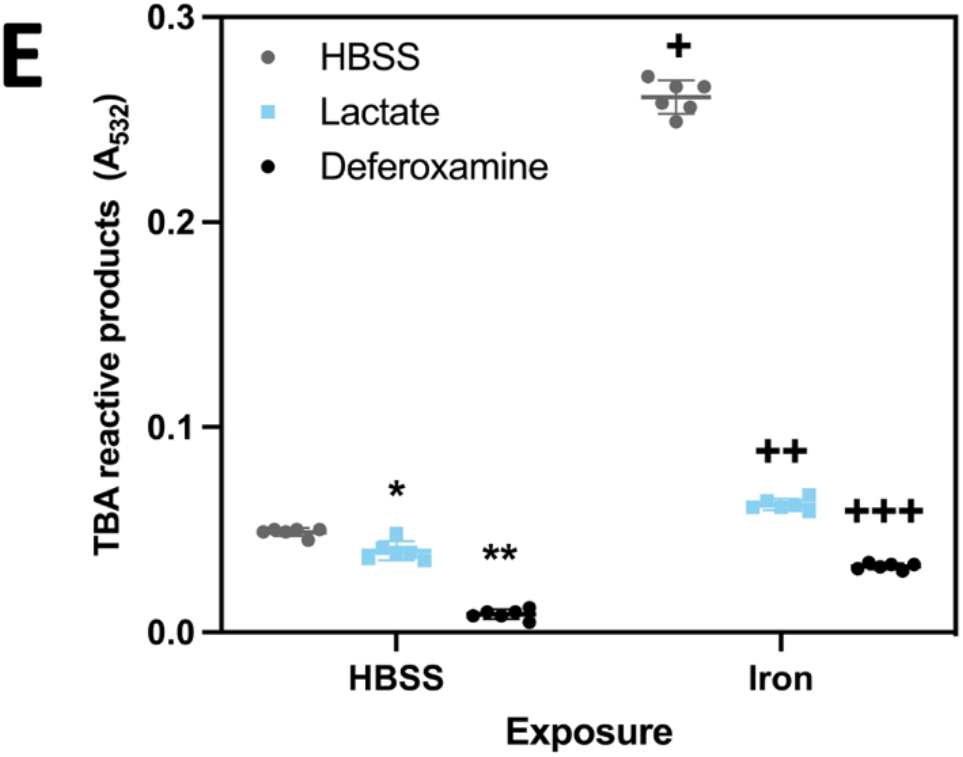
Fe3+ increased while inclusion of lactate and deferoxamine significantly decreased thiobarbituric acid (TBA) reactive products within an acellular incubation. *= significantly decreased relative to HBSS, **= significantly decreased relative to other HBSS exposures, +significantly increased relative to HBSS incubation, ++= significantly decreased relative to exposures, +++= significantly decreased relative to other Fe3+ exposures, p<0.05.

### Cell lactate dehydrogenase (LDH) and lactate decrease with exposure to ferric ammonium citrate (FAC)

Associations of both LDH and lactate with iron available were evaluated in cells. RT-PCR revealed a lack of a significant elevation in both HIF-α mRNA/β-Actin mRNA and LDH mRNA/β-Actin mRNA after 24-hour incubation of Caco-2 cells with 50 μM deferoxamine (**Table 1**). However, cell exposure to 500 μM FAC for 24 hours decreased HIF-α mRNA/β-Actin mRNA and LDH mRNA/β-Actin mRNA (**Table 1**). LDH concentrations in Caco-2 cell scrapings did not change with time after exposures to HBSS (5A: 56.333 units/L ± 10.328 at 4 hours, 61.167 units/L ± 14.919 at 8 hours, 66.167 units/L ± 13.644 at 24 hours) (5B: 65.167 units/L ± 13.717 at 4 hours, 60.167 units/L ± 13.288 p = 0.8022 at 8 hours, 65.500 units/L ± 13.766 at 24 hours) (**Figure 6A and 6B**). While exposure to the iron chelator deferoxamine did not impact cell LDH levels, incubations with FAC significantly decreased concentrations of this enzyme at 8 and 24 hours (54.500 units/L ± 9.935 at 4 hours, 58.333 units/L ± 10.013 p = 0.7752 at 8 hours, 53.333 units/L ± 9.070 p = 0.9763 at 24 hours with 25 μM deferoxamine; 51.333 units/L ± 9.564 at 4 hours, 49.667 units/L ± 9.750 at 8 hours, 48.500 units/L ± 8.503 p = 0.8588 at 24 hours with 50 μM deferoxamine; 65.167 units/L ± 10.284 at 4 hours, 48.167 units/L ± 9.283 at 8 hours, 26.667 units/L ± 7.202 at 24 hours for FAC 100 μM, 58.167 units/L ± 10.647, 28.333 units/L ± 6.772, 22.667 units/L ± 6.282 p < 0.0001 for FAC 500 μM) (**Figure 6A and 6B**). Following exposures of Caco-2 cells to HBSS, lactate levels in both the cell scrapings and supernatant increased with time of incubation (6A: 0.540 mg/dL ± 0.087 at 4 hours, 0.639 mg/dL ± 0.079 at 8 hours, 1.749 mg/dL ± 0.238 at 24 hours) (6B: 0.553 mg/dL ± 0.113 at 4 hours, 0.798 mg/dL ± 0.131 at 8 hours, 2.000 mg/dL ± 0.309 at 24 hours) (6C: 19.550 mg/dL ± 3.012 at 4 hours, 23.900 mg/dL ± 1.880 at 8 hours, 31.933 mg/dL ± 4.385 at 24 hours) (6D: 19.517 mg/dL ± 2.495 at 4 hours, 23.917 mg/dL ± 3.456 at 8 hours, 33.567 mg/dL ± 4.814 at 24 hours) (**Figure 7A-D**). Exposures to deferoxamine did not show an impact on either cell or supernatant lactate concentrations (7A: 0.581 mg/dL ± 0.082 at 4 hours, 0.901 mg/dL ± 0.142 at 8 hours, 0.950 mg/dL ± 0.159 when compared to concentration at 8 hours at 24 hours for deferoxamine 25μM; 0.609 mg/dL ± 0.101 at 4 hours, 1.171 mg/dL ± 0.161 p < 0.0001 at 8 hours, 0.949 mg/dL ± 0.174 when compared to concentration at 8 hours at 24 hours for deferoxamine 50μM) (7C: 18.467 mg/dL ± 2.857 at 4 hours, 23.900 mg/dL ± 3.741 at 8 hours, 21.700 mg/dL ± 2.234 at 24 hours for deferoxamine 25μM, 31.900 mg/dL ± 4.556 at 4 hours, 30.317 mg/dL ± 4.665 at 8 hours, 32.217 mg/dL ± 3.848 at 24 hours for deferoxamine 50μM with). Incubations with FAC did reveal decreased lactate levels in both cell scrapings and supernatant (7B: 2.000 mg/dL ± 0.309 with HBSS at 24 hours 1.643 mg/dL ± 0.170 at 24 hours with FAC 100μM, 0.965 mg/dL ± 0.150 at 24 hours for FAC 500μM) (7D: 33.567 mg/dL ± 4.814 with HBSS at 24 hours, 24.717 mg/dL ± 3.342 with FAC 100μM at 24 hours, 22.900 mg/dL ± 4.208 with FAC 500μM at 24 hours). These decrements in lactate concentrations in the cell lysates and supernatants were evident at 4, 8, and 24 hr.

**Table 1:** RT-PCR for HIF-1α and lactate dehydrogenase reveals significant relationship between FAC and HIF-α mRNA/β-Actin mRNA and LDH mRNA/β-Actin mRNA.

| Exposure | HIF-1α mRNA/β-Actin mRNA | LDH mRNA/β-Actin mRNA |
| --- | --- | --- |
| HBSS | 0.68 ± 0.18 | 0.41 ± 0.11 |
| Deferoxamine 50 μM | 0.72 ± 0.28 | 0.46 ± 0.11 |
| Ferric ammonium citrate 500 μM | 0.21 ± 0.12* | 0.20 ± 0.10* |

\*Significantly different relative to HBSS exposure, $p < 0.05$ , standard deviation shown.

**Figure 6.**
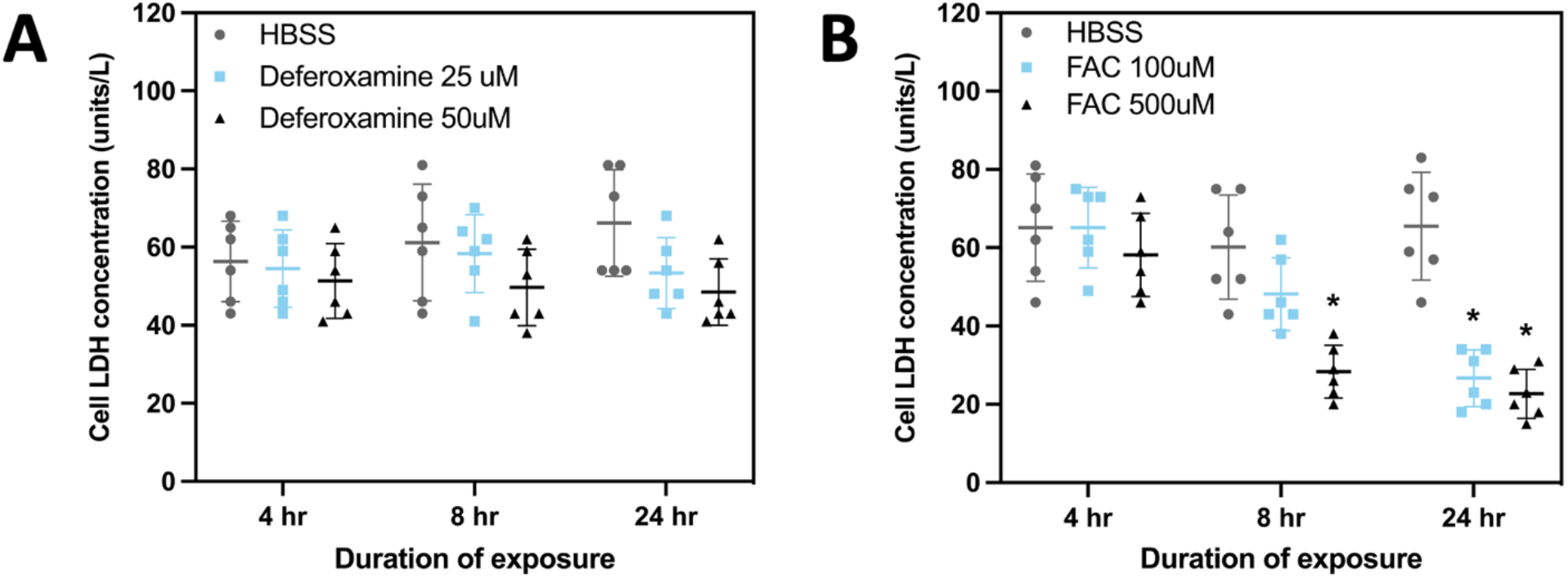
Cell LDH concentrations. ANOVAs demonstrated differences between [LDH] in Caco-2 (A and B) after exposure to HBSS, deferoxamine, and FAC for 4, 8, and 24 hr. *= Significantly different relative to HBSS with p<0.05.

**Figure 7.**
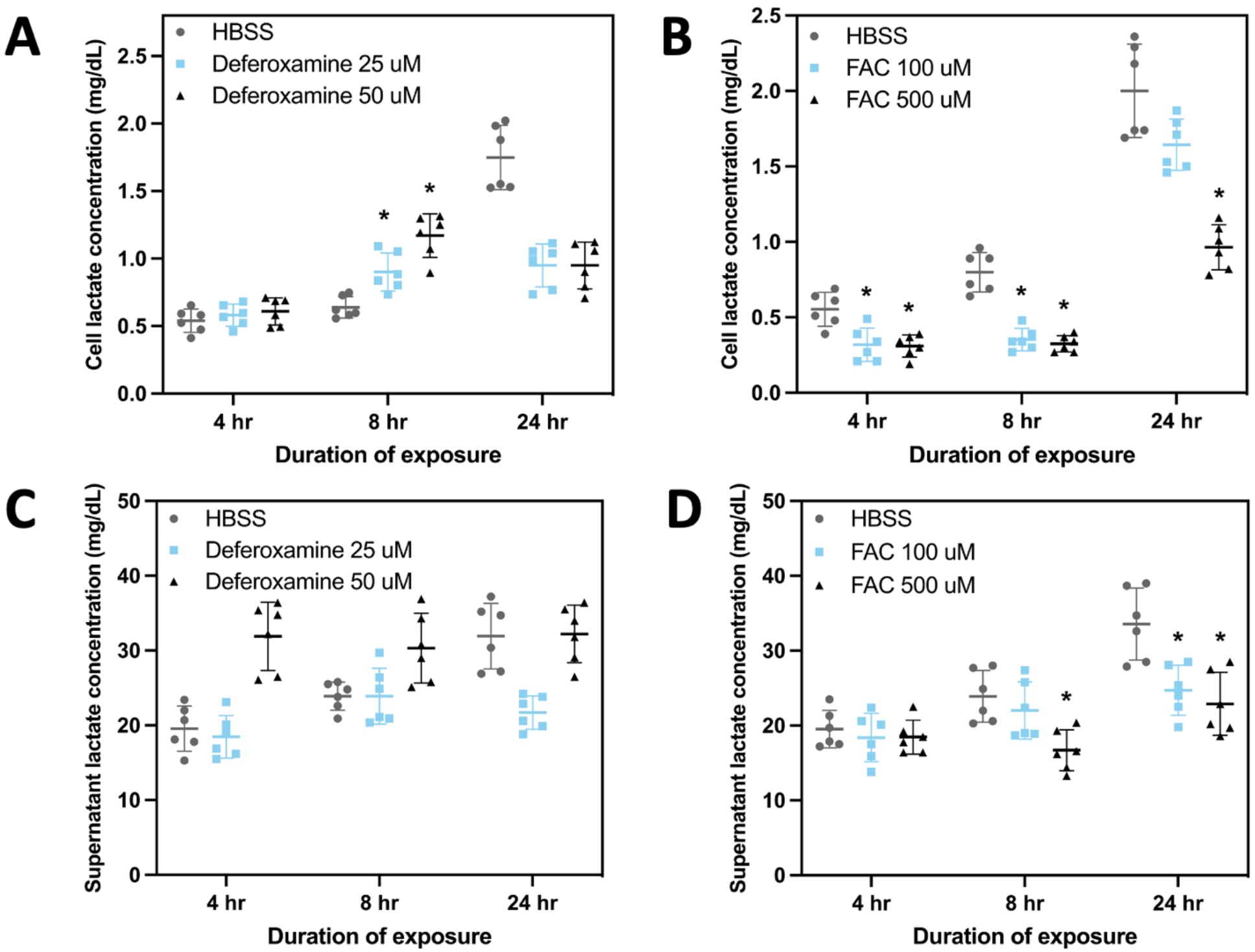
Cell and supernatant lactate concentrations indicate decreases in cell lactate concentration after exposure to FAC. ANOVAs demonstrated differences between lactate concentrations in Caco-2 cell scrapings (A and B) and supernatants (C, D) after exposure to HBSS, deferoxamine, and FAC for 4, 8, and 24 hr. *= significantly different relative to HBSS with p<0.05.

**Figure 8.**
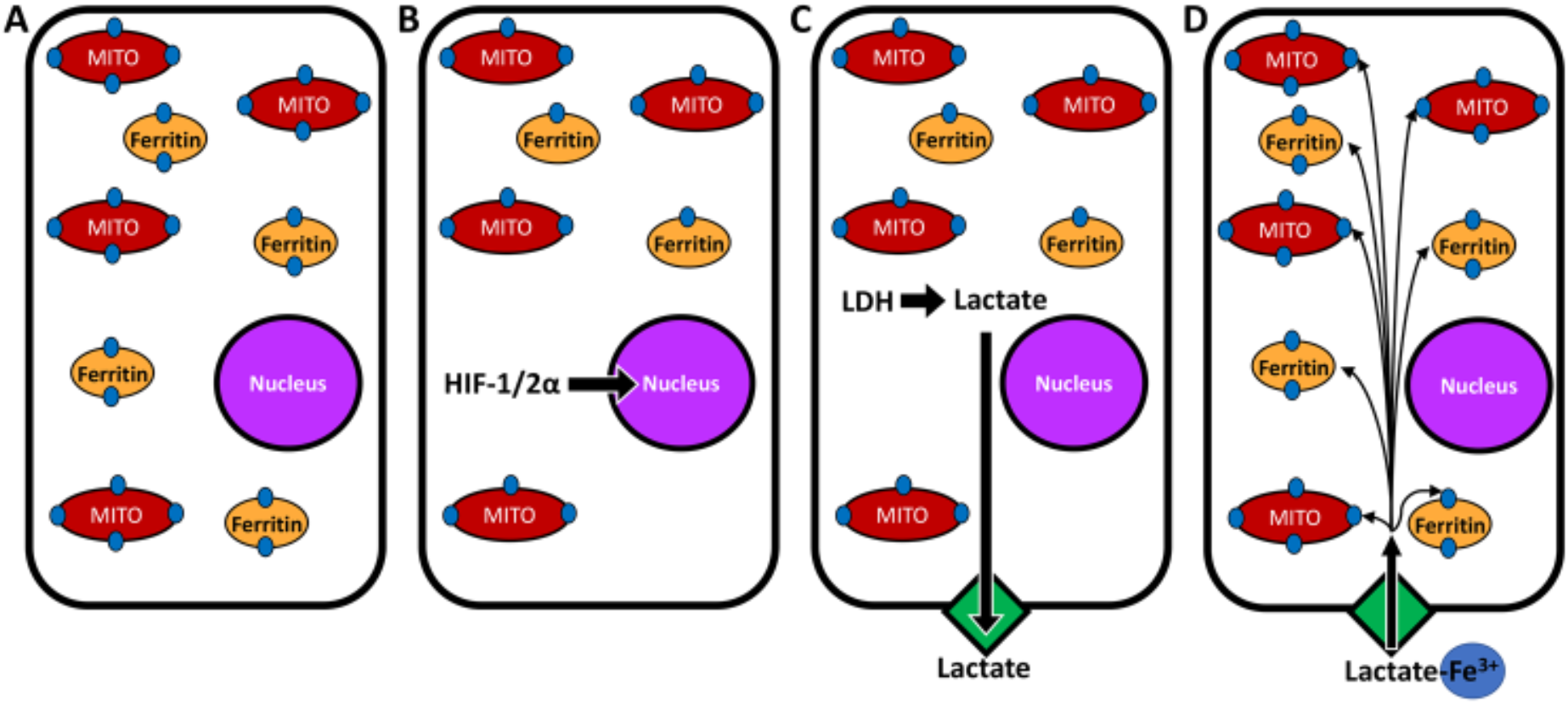
Schematic for changes in iron homeostasis, lactate generation, and import. Cell iron homeostasis (A) is disrupted with absolute or functional iron deficiency. In response to a reduction in intracellular iron, HIF-1/2α is activated and this transcription factor moves to the nucleus to impact the expression of numerous proteins including lactate dehydrogenase (B). Lactate generation increases and is released from the cell (C). After the complexation of available iron, the lactate binds with receptors and delivers iron to the cell reversing the absolute or functional metal deficiency (D).

### Associations between LDH and iron homeostasis demonstrate an inverse relationship between LDH and indices of iron homeostasis

The National Health and Nutrition Examination Survey (NHANES) was used to investigate the relationship between LDH concentrations and indices of iron homeostasis in non-pregnant females, age 20-49. The cohort from the NHANES (n= 621) had a mean age of 34.2 ± 8.4 yr (range of 20-49 yr). This included Hispanic-American (44.4%), non-Hispanic whites (32.4%), non-Hispanic blacks (18.4%), and other (4.8%) females. Mean (± std. dev.) LDH concentration and indices of iron homeostasis derived from data from NHANES are provided (**Table 2A**). Spearman correlations demonstrated significant negative relationships between the lactate dehydrogenase (LDH) concentration and several indices of iron homeostasis include serum iron, mean corpuscular hemoglobin (MCH), and mean corpuscular volume (MCV) (**Table 2B**).

**Table 2A.** Mean (± std. dev.) for lactate dehydrogenase and indices of iron homeostasis in the NHANES 2005-2010 study cohorts.

|  | **Mean ± std. dev.** | **Range** |
| --- | --- | --- |
| **LDH (U/L)** | 123.8 ± 26.6 | 51-341 |
| **Serum iron (μg/dL)** | 73.7 ± 37.4 | 7-382 |
| **Serum ferritin (ng/mL)** | 46.4 ± 45.6 | 2-300 |
| **Hemoglobin (g/dL)** | 13.1 ± 1.2 | 6.2-16.2 |
| **Mean cell hemoglobin (pg)** | 29.6 ± 2.6 | 14.7-37.0 |
| **Mean corpuscular volume (fL)** | 86.9 ± 6.3 | 51.1-104.4 |

**Table 2B.** Spearman correlations reveal significant negative relationships between lactate dehydrogenase concentration and indices of iron

|  | **Correlation** | **p value** |
| --- | --- | --- |
| **Serum iron (µg/dL)** | -0.123 | 0.002 |
| **Serum ferritin (ng/mL)** | +0.081 | 0.065 |
| **Hemoglobin (g/dL)** | -0.015 | 0.71 |
| **Mean cell hemoglobin (pg)** | -0.112 | 0.005 |
| **Mean corpuscular volume (fL)** | -0.104 | 0.009 |

## Discussion

### Lactate increases cell iron uptake

Relative to both other metal lactates and iron complexed to citrate, Caco-2 intestinal epithelial cells demonstrated a greater capacity to import metal from iron lactate. Exposure of Caco-2 and BEAS-2B lung epithelial cells to Na lactate and ferric ammonium citrate (FAC) resulted in a greater iron import compared with incubations with FAC alone. With a pKa approximating 3.9, the inclusion of lactic acid in cell incubations was anticipated to be equivalent to that of Na lactate and it similarly elevated non-heme iron concentration following incubations with FAC. These results are comparable to those of prior studies which demonstrated that lactate could increase iron import by Caco-2 cells ^25^. Further, this α-hydroxycarboxylate elevates iron uptake with findings that lactic fermentation of foods increased metal availability ^26,27^. Exposure to other organic acids had an equivalent capacity to increase iron import into cells. Previous investigation of exposures to citrate, fumarate, malate, oxalate, succinate, and tartrate demonstrated that carboxylates other than lactate can impact iron homeostasis ^10^. Among these, the α-hydroxycarboxylates appeared to have an increased ability to participate in iron import.

Prior work suggests that the alcohol functional group in α-hydroxycarboxylates deprotonates and participates in complex formation with iron ^9^. Among carboxylates, citrate is commonly utilized by plants and microbes in iron transport ^9^. The complexation of iron by citric acid involves coordination both by the carboxylate and hydroxyl groups. Citrate solubilizes iron and increases bioavailability in plants and microbes. Subsequently, iron-citrates are a constituent of the extracellular and cytoplasmic low molecular iron pools in vertebrates as well as plants. Our study supports lactate functioning in a capacity comparable to citrate, and other carboxylate/hydroxycarboxylate siderophores, with iron complexation by the organic acid followed by cell metal uptake and increased availability resulting. Prior work supports a capacity of lactate to complex/chelate iron with microbial exposure to high concentrations upregulating genes involved in the homeostasis of this metal ^14^. Increased iron import was correlated with increasing levels of the storage protein ferritin and the concentration of metal associated with mitochondria. This may contrast prior investigation which suggested that lactate addition to Caco-2 cells did not affect ferritin formation following co-incubation with iron ^27^. The role of lactate in cell iron transport, rather than another carboxylate/hydroxycarboxylate, is likely associated with its higher concentrations in both cells and extracellular fluids (e.g. human blood lactate is 1 to 2 mM but can be elevated to levels as high as 25 mM) in addition to its ability to cross the cell membrane using a monocarboxylate transporter ^28^.

There is conflicting data regarding iron import following exposures to compounds/substances which can complex/chelate iron, including lactate, with many being reported to also block cell iron import ^29–31^. Inconsistencies can result from several factors including 1) the magnitude of the stability constant of the coordination complex with the formation of a strong chelate preventing cell import of iron, 2) the type and solubility of the coordination complex, 3) pH value in the medium, cell, and lysosome, 4) chemical reduction and oxidation of iron cation by the compound/substance which complexes/chelates, 5) affinity for an uptake system and ability to deliver the metal to uptake proteins, 6) concentrations of both the compound/substance and iron, and 7) whether the metal is ferric or ferrous ^9,10,32^. Accordingly, investigations show a limited range of molar ratios in which these compounds/substances enhance iron uptake and the range varies by each compound with an increase in the molar ratio resulting in loss of promotion and ultimately inhibition of iron uptake.

### Cell lactate dehydrogenase (LDH) and lactate correspond with iron availability

LDH expression in Caco-2 cells was associated with iron availability. Both LDH expression and activity were previously shown to be increased by hypoxia ^33,34^. In many cell types, this elevation in LDH expression with hypoxia was regulated transcriptionally by HIF-1/2α ^35,36^. Investigations have repeatedly demonstrated that, in addition to being controlled by hypoxia, HIF-1/2α is regulated by the level of available iron ^37–39^. Because of its chemical properties, iron is singular among the metals in its capacity to support biochemical reactions of oxygen ^40^. Accordingly, significant overlap exists in the regulation of oxygen and iron homeostasis; the homeostasis of each is regulated by the other ^41^. The oxygen dependency of prolyl-hydroxylase domain enzymes (PHDs) which target HIF for proteasomal degradation provides the basis of a widespread oxygen-sensing mechanism ^42^. PHDs are also iron-dependent and metal deficiency will stabilize this transcriptional factor. Subsequently, HIF-1/2α not only regulates the expression of proteins pivotal in oxygen metabolism but also many essential proteins in iron homeostasis (e.g. divalent metal transporter 1, ferroportin, transferrin receptor, hepcidin, and heme oxygenase) ^43,44^. The results of this study suggest that LDH is another example of a protein required for iron homeostasis whose expression is controlled by HIF-1/2α.

Cell LDH activity and lactate concentrations both corresponded to metal availability with decrements following exposure to higher levels of iron. It is unclear why the expression of LDH and lactate did not reproducibly increase with deferoxamine which can impact a cell iron deficiency but may reflect early cell demise anticipated with the use of the chelator. These results do support a relationship of LDH expression and/or activity to iron homeostasis with decrements following exposures to iron compounds. This is comparable to a previous study of primary human cardiac myocytes in which LDH activity was decreased by iron supplementation ^45^. LDH expression and activity in *Neisseria* gonorrhoeae and *Mycobacterium* smegmatis similarly increased with iron-limitation ^46,47^. The inclusion of an iron chelator (e.g. 2,2’-bipyridine or 1,10-phenanthroline) in the culture of mammalian cells also increased the synthesis of LDH ^48,49^.

### Associations between LDH and blood indices of iron homeostasis

Concentrations of LDH in blood showed an inverse relationship with iron, MCH, and MCV in the NHANES 2005-2010 cohort ^22–24^. This supports a possible role for lactate, the LDH product, in the provision of iron to cells. In animals, both dietary restriction of iron and its resultant iron-deficiency anemia were found to elevate the total LDH activity in muscles and an accumulation of lactate ^50,51^. Absolute iron deficiency in humans, frequently associated with anemias, has been repeatedly associated with elevations in lactate ^52,53^. Increased iron availability can reverse these elevations in lactate ^54^. Transfusion of packed erythrocytes or washed erythrocytes to patients with megaloblastic anemia, which increased iron availability, invariably resulted in a decline of serum LDH ^55^. Equivalently, exposures that impact a functional iron deficiency (e.g. numerous infections and particles) have previously been shown to affect elevations in blood LDH and lactate ^56,57^. Hemoglobin (HGB) and ferritin showed no significant associations with LDH. The lack of significant correlations with LDH can reflect 1) the failure of HGB to change with other indices of iron deficiency and 2) the impact of inflammation on ferritin ^58–60^.

Cell iron deficiency is predicted to contribute to increased LDH expression/activity and subsequent hyperlactatemia and lactic acidosis, two syndromes associated with human morbidity and mortality. Many of these presentations can be classified as Type B lactic acidosis (lactic acidosis without evidence of systemic hypoperfusion). Among the causes of Type B lactic acidosis (medications, toxic exposures, diabetes mellitus, alcoholism, HIV infection, and mitochondrial dysfunction) are numerous associations with iron deficiency (e.g. exposures to catechols which are metal chelators, and mitochondrial encephalomyopathy with lactic acidosis and stroke-like episodes syndrome) ^61,62^. The association between iron availability and lactatemia/lactic acidosis would allow understanding the relationships between concentrations of this α-hydroxycarboxylate in specific tissues deficient in iron as well as with anemias and their resolution ^52,54,63,64^. Finally, it is also recognized that LDH participates in tumor initiation and growth and development of metastases with overexpression decreasing survival among cancer patients ^65,66^. These relationships possibly reflect the participation of LDH and lactate in iron homeostasis in cancer cells ^67,68^.

## Conclusion

*In vitro* models are paramount to understanding of iron import and homeostasis in human cells. We have identified that lactate may play a role in the management of iron in human tissues comparable to plant cells and microbes. Cell lactate dehydrogenase (LDH) activity, which is responsible for lactate production, reflected iron availability in our human epithelial cultures. Blood concentrations of LDH in a human cohort correlated with indices of iron homeostasis further supporting a possible role for lactate in the transport of the metal. Future work must investigate the genetic manipulation of LDH, and subsequent lactate levels, with observations of the impact on iron homeostasis.

## Acknowledgements

The authors thank the Department of Chemical Engineering for startup funds (AK, RK) as well as the Office of the Provost PEAK awards (CG).

## Author Contributions

C.G., A.N.K., and A.G. conceived the study. C.G., A.N.K., A.G., and R.A.K. provided experimental design input. C.G. conducted experiments with Caco-2 cells and was the primary manuscript author. A.G, J.S., L.D. conducted experiments with BEAS-2B cells. C.G. and A.G. analyzed experimental data. D.S. analyzed NHANES data. All authors provided input toward the manuscript.

## Data availability

The datasets generated and analyzed during the current study are available from the corresponding author on reasonable request.

